# Bootstrap Distillation: Non-parametric Internal Validation of GWAS Results by Subgroup Resampling

**DOI:** 10.1101/104497

**Authors:** David A. Eccles, Rodney A. Lea, Geoffrey K. Chambers

## Abstract

Genome-wide Association Studies are carried out on a large number of genetic variants in a large number of people, allowing the detection of small genetic effects that are associated with a trait. Natural variation of genotypes within populations means that any particular sample from the population may not represent the true genotype frequencies within that population. This may lead to the observation of marker-disease associations when no such association exists.

A bootstrap population sub-sampling technique can reduce the influence of allele frequency variation in producing false-positive results for particular samplings of the population. In order to utilise bioinformatics in the service of a serious disease, this sub-sampling method has been applied to the Type 1 Diabetes dataset from the Wellcome Trust Case Control Consortium in order to evaluate its effectiveness.

While previous literature on Type 1 Diabetes has identified some DNA variants that are associated with the disease, these variants are not informative for distinguishing between disease cases and controls using genetic information alone (AUC=0.7284). Population sub-sampling filtered out noise from genome-wide association data, and increased the chance of finding useful associative signals. Subsequent filtering based on marker linkage and testing of marker sets of different sizes produced a 5-SNP signature set of markers for Type 1 Diabetes. The group-specific markers used in this set, primarily from the HLA region on chromosome 6, are considerably more informative than previously known associated variants for predicting T1D phenotype from genetic data (AUC=0.8395). Given this predictive quality, the signature set may be useful alone as a screening test, and would be particularly useful in combination with other clinical cofactors for Type 1 Diabetes risk.

## 1 Background

Personalised medical treatment based on genome profiles is a major goal of genetic research in the 21^*st*^ century [see 2, 10]. However, complex genotype-environment interactions for common diseases make it difficult to determine which specific genetic features should be used to construct such profiles. Hence the prediction of genetic risk is a major challenge of the 21^*st*^ century.

The introduction of large-scale Single Nucleotide Polymorphism (SNP) genotyping systems has enabled genetic variants to be typed *en-masse*, shifting the main effort required in a genetic risk study from genotyping to data analysis (or bioinformatics). Here we investigate genetic markers for Type 1 Diabetes (T1D), demonstrating how a population sub-sampling method may assist in the identification of risk markers for a complex disease.

### 1.1 Type 1 Diabetes

Type 1 Diabetes mellitus (T1D) is a disorder typically characterised by an absence of insulin-producing beta cells in the pancreas, either through loss of the cells themselves, or through the reduction in capacity of the cells to produce insulin [see 1]. This disorder shares with the more common Type 2 Diabetes mellitus (T2D) a characteristic symptom of high blood glucose levels. In some cases, this glucose also passes through to the urine, creating a sticky/sweet substance that attracts ants [see 5, pp. 7,11]. In T2D, this high blood glucose is caused by cells not responding to insulin (insulin resistance), while in T1D the excess is caused by a reduction in insulin production (insulin dependence).

The incidence of T1D varies throughout the world, with rates of incidence as low as 0.0006% per year in China, 0.02% in the UK, up to nearly 0.05% per year in Finland. About 50-60% of cases of T1D manifest in childhood (younger than 18 years), and the disease is believed to be caused by an abnormal immune response after exposure to environmental triggers such as viruses, toxins or food [see 3]. While a spring birth is correlated with T1D risk, the *diagnosis* of Type 1 Diabetes is more common in autumn and winter [see 1].

#### 1.1.1 Symptoms, Diagnosis and Management of T1D

Typical symptoms of T1D include excess urine output (polyuria), thirst and increased fluid intake (polydypsia),blurred vision, and weight loss. When left untreated, this form of diabetes can lead to a build-up of ketone bodies and a reduction of blood pH (ketoacidosis), reducing mental faculties and causing a loss of consciousness [see 5, p. 7].

Diabetes can be diagnosed by a single *random*^1^ blood glucose test, as long as symptoms are present and blood glucose levels are found to be in excess (typically *>* 11.1 *mmol l^−^*^1^) of those normally observed. In situations where symptoms are less obvious and/or glucose levels are at the high end of the normal range, a glucose tolerance test (GTT) is used for diagnosis. In this test, fasting patients have their blood glucose level tested, patients then consume a measured dose of oral glucose, and blood glucose levels are measured 2 hours later. A fasting glucose level in excess of 6.1 *mmol l^−^*^1^, or post-load level in excess of 11.1 *mmol l^−^*^1^ is considered diagnostic for both forms of Diabetes Mellitus. Type 1 Diabetes (as distinct from T2D) encompasses a range of diseases that involve autoimmunity. It can be diagnosed by the presence of antibodies to glutamic acid decarboxylase, islet cells, insulin, or ICA512 [see 5, p. 19].

As the symptoms of T1D are caused by high blood glucose levels (hyperglycaemia) due to a lack of insulin, these symptoms can be relieved by the introduction of insulin into the blood. This is typically carried out by supplying measured doses of insulin via intramuscular injections or by the use of insulin pumps [see 3]. Individuals with T1D need a constant supply of insulin for survival, together with occasional insulin bursts to control variable blood glucose levels throughout the day (e.g. after meals). In contrast, individuals with T2D only require insulin for survival in rare cases [see 5, p. 16]. Slow-release insulin and consumption of foods with a low glycaemic index can help to reduce the extremes of T1D symptoms.

Improperly managed treatment can cause further medical complications in a diabetic patient. Too much insulin, excessive physical activity, or not enough dietary sugar can result in low blood glucose levels (hypoglycaemia), which produce short-term autonomic and neurological problems such as trembling, dizziness, blurred vision, and difficulty concentrating. Hypoglycaemia is treated either by ingestion of sugar, or by intravenous glucose in severe cases [see 3].

#### 1.1.2 Complications of T1D

The initial symptoms of T1D are not usually severe, and the disease may progress for a few years before a diagnosis is made and treatment is given. However, long-term complications can appear when the disease is not managed appropriately [see 5, p. 8]. Retinal damage progresses in about 20-25% of individuals with T1D, with later stages causing retinal detachment and consequent loss of sight. Renal failure is also a problem in diabetic individuals, which is indicated by high urinary protein levels. When individuals have these high levels, progression to end-stage renal disease occurs in about 50% of cases. Neural defects are also a potential complication of T1D, most commonly damage to peripheral nerves, leading to ulceration, poor healing and gangrene unless good care is taken of the body extremities [see 3].

#### 1.1.3 Genetic Contribution to T1D Risk

Type 1 Diabetes has a heritability of around 88% [6], indicating that a substantial proportion of variance in disease susceptibility can be attributed to genetic factors. About 50% of the genetic contribution to T1D can be accounted for by variation in the HLA region on chromosome 6, and 15% is accounted for by variation in two other genes, IDDM2 and IDDM12 [see 3]. Incidence rates in migrant populations quickly converge to those of the background population, suggesting that although the genetic contribution to the disease is high, environmental factors probably play a significant role in triggering the onset of disease [see 3].

### 1.2 Wellcome Trust Case Control Consortium Study

The Wellcome Trust Case Control Consortium (WTCCC, http://www.wtccc.org.uk) was established in 2005 to identify novel genetic variants associated with seven common diseases, including Type 1 Diabetes [12]. 2000 individuals with T1D, and 1500 individuals from the National Blood Service (NBS)^2^ were genotyped for the WTCCC using an Affymetrix GeneChip 500k Mapping Array Set.

The Wellcome Trust Case Control Consortium [12] reported associations near five gene regions that had been previously associated with T1D: The major histocompatibility complex (MHC) on chromosome 6, CTLA4 and IFIH1 on chromosome 2, PTPN22 on chromosome 1, and IL2RA on chromosome 10. The insulin gene (INS) on chromosome 11 was also associated with T1D; the only SNP tagging INS failed quality control filters, but also indicated strong association with T1D when examined. A number of other regions showed evidence of association with T1D in the Wellcome Trust Case Control Consortium [12] study: 4q27 (chromosome 4); 10p15 (chromosome 10); 12p13, 12q13 and 12q24 (chromosome 12) 16p13 (chromosome 16); and 18p11 (chromosome 18). Most of these regions include genes involved in the immune system. However, only two genes are in 16p13, and both have unknown functions (KIAA0350 and dexamethasone-induced transcript). The strongest association signal for T1D was detected within the HLA region of chromosome 6, a region in which multiple SNPs had strong associations with T1D, but only one of those SNPs (rs9272346) was reported in the results table of the strongest associations [see 12, table 3].

### 1.3 Replication Issues in GWAS

The Genome-wide Association Study (GWAS) is a common method for discovering genetic contributions to complex human diseases. The outcome of these studies is to determine the degree of association between single genetic markers and a heritable trait. Commonly, an analysis is carried out on a large number of genetic variants in a large number of people, allowing the detection of small genetic effects that are associated with a trait. In recent years, an initial search for variants is carried out by whole-genome sequencing in a small sub-population to identify variants that are common in the population of interest.

A study style that is built around correlation and association rather than a hunt for causal variants requires extreme care to ensure that observed associations are valid *and* causal. Studies need to have good within-study validation to reduce the likelihood of false-positive results being obtained and treated as true associations, and need to be supported by good independent validation. The distinction between association and causation is important – GWAS are used as hypothesis-generating tools to narrow down, through association, the search for potential causative loci. After the associations have been validated, it is expected that they will be followed up with studies attempting to determine the true causative status of that association.

Such causative studies are difficult, and progress towards understanding the aetiology of common disease has been slow [see 4].

### 1.4 Sampling Errors in GWAS

Natural variation of genotypes within populations means that any particular sample from the population may not represent the true genotype frequencies within that population. This may lead to the observation of marker-disease associations when no such association exists. This is particularly important when considering populations with mixed ancestry, where markers that are informative for distinguishing population ancestry may become accidentally associated with a particular disease [see 9].

Bootstrapping by repeated re-sampling of a representative draw made from a group can estimate population variation in genotype frequencies by observing variation within the sub-samples. A re-sampling technique, as presented here, can reduce the influence of allele frequency variation by excluding false-positive results that are specific for particular samplings of the population.

## 2 Method and Results

### 2.1 Method Summary

The Wellcome Trust Case Control Consortium (WTCCC) have genotyped 2000 individuals diagnosed with T1D, and 1500 individuals from the National Blood service (NBS) using the Affymetrix 500k chip (500568 SNPs). These genotypes were obtained from WTCCC for subsequent computer-based research exploring the utility of the author’s new bootstrap sub-sampling method for genome-wide association studies. Genotype data were filtered at a SNP level to remove those SNPs that were present on the X chromosome; individuals flagged by WTCCC as having potentially invalid genotype data were removed from the dataset.

The study group was split into two equal-sized groups: a *discovery* group (981 T1D cases, 729 NBS controls), and a *validation* group (982 T1D cases, 729 NBS controls). Subsequent filtering, analysis, and selection of SNPs was carried out on the discovery group, while the validation group was *only* used to test the effectiveness of the selected SNP set in a situation distinct from that used to generate this set of SNPs (see Figure 1).

A bootstrap sub-sampling method was used to reduce the initial panel of 500k SNPs down to a set that consistently produced associations on all bootstrap sub-samples. Sub-samples of the Type 1 Diabetes (T1D) cases and National Blood Service (NBS) controls were used to estimate the in-group variance of association statistics throughout the genome. Markers that were informative and had low variance were selected as candidate markers for a minimal informative set of 45 markers.

The final refinement step tested sets of SNPs in combination, rather than single SNPs alone, in the hope that those sets would be able to capture a wider range of genetic variation than any single marker (or combination of data for single markers alone) could provide. Once a suitable SNP set was found, that set was tested in the validation group to confirm the utility of the set for distinguishing T1D cases from NBS controls.

### 2.2 Genotyping and Filtering of Individuals and SNPs

Genotype data from 2000 T1D cases and 1500 NBS controls were provided by WTCCC. This genotyping had been carried out on an Affymetrix 500k SNPchip (500568 SNPs). The purpose of the initial genotyping procedure is to obtain a large sample of candidate markers (*>* 100, 000) from which to pick the most informative.

Due to sex differences in expression for X-chromosome SNPs, all 10536 X chromosome SNPs were excluded. The WTCCC dataset also included a list of 37 T1D cases and 42 NBS controls to exclude for a number of reasons (e.g. high proportion of missing genotype data, duplicate individuals, non-European ancestry), so these individuals were also removed from the present study (X chromosome filtered set of 490032 SNPs, 1963 cases, 1458 controls).

**Figure 1:**
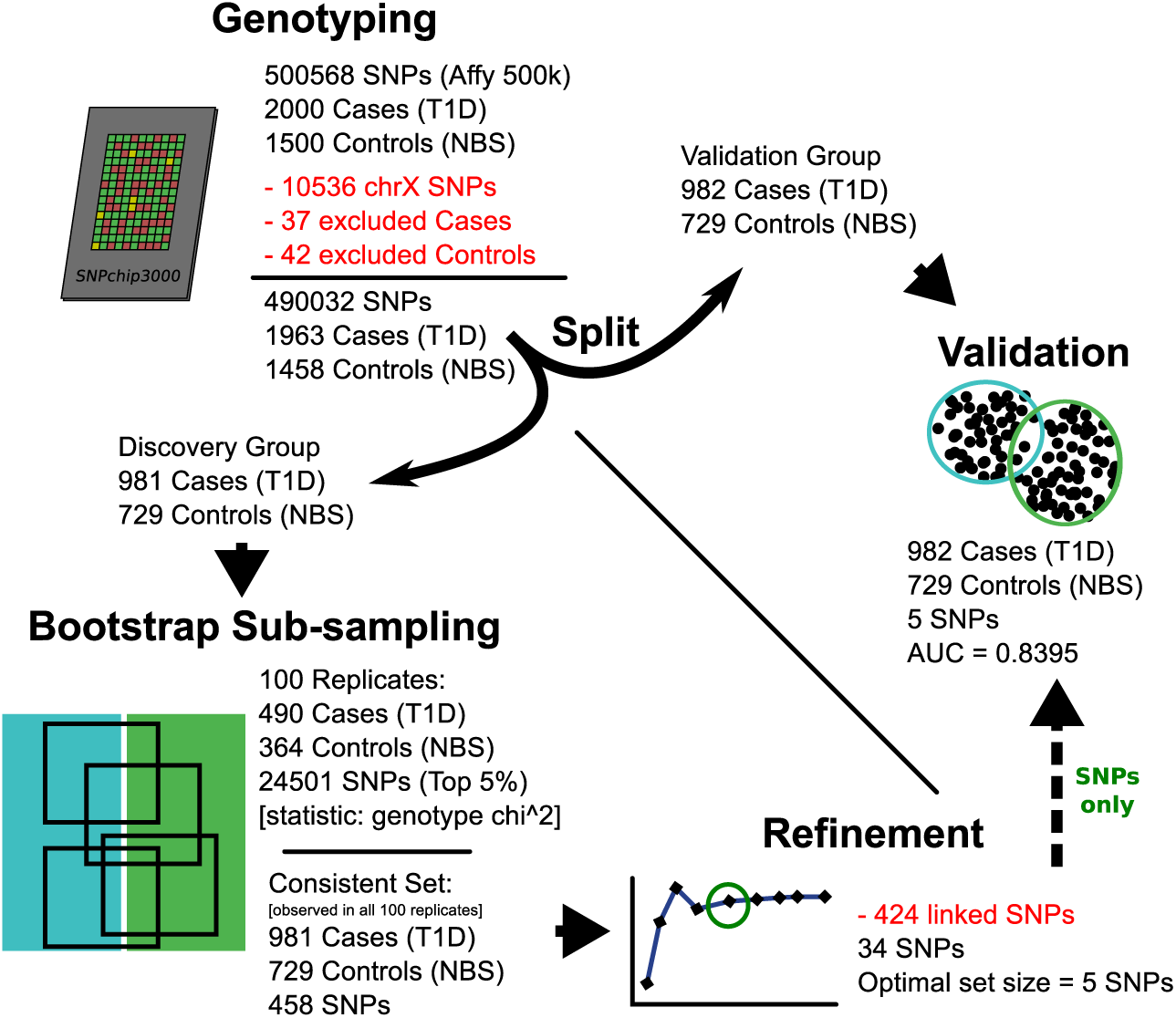
An overview of the marker set construction procedure, using an initial validation/discovery split, bootstrap sub-sampling, set refinement, and internal validation.

#### 2.2.1 Separation of Discovery and Validation Groups

Individuals were randomly assigned into one of two groups: a discovery group with 981 T1D cases and 729 NBS controls, and a validation group with 982 T1D cases and 729 NBS controls. To ensure a robust analysis, the validation group was only used for the final validation of a generated SNP set, and not used for any part of the SNP discovery procedures.

#### 2.2.2 Marker Association Values Across the Entire Genome

Association scores were calculated across the entire autosomal genome (see Figure 2). A genotype *χ*^2^ test was used, comparing observed genotype counts for each group to expected genotype counts for both groups combined. High association scores (*χ*^2^ *>* 100) were found on chromosomes 3, 6, 12, 16, and 22, but the scores were only consistently high for a region of about 10Mb in the middle of the short arm of chromosome 6 (30-40Mb from the 5' end of the reference strand).

### 2.3 Bootstrap Sub-sampling of the Discovery Group

A bootstrap sub-sampling method was then carried out, generating 100 subsample replicates of the discovery group, each replicate having 490 T1D cases and 364 NBS controls (i.e. retaining the same proportions as the original 981 cases and 729 controls), sampled from the original discovery set without replacement. The SNPs were then ranked by *χ*^2^, and a *bootstrap-consistent set* of 458 SNPs was identified, each ranked in the top 5% of SNPs (24501 SNPs) in every bootstrap sub-sample (see Figure 3-A). Most of these 458 SNPs had a maximum rank below 5000, whereas most of the remaining 489574 SNPs had a maximum rank above 350000 (see Figure 3-B).

**Figure 2:**
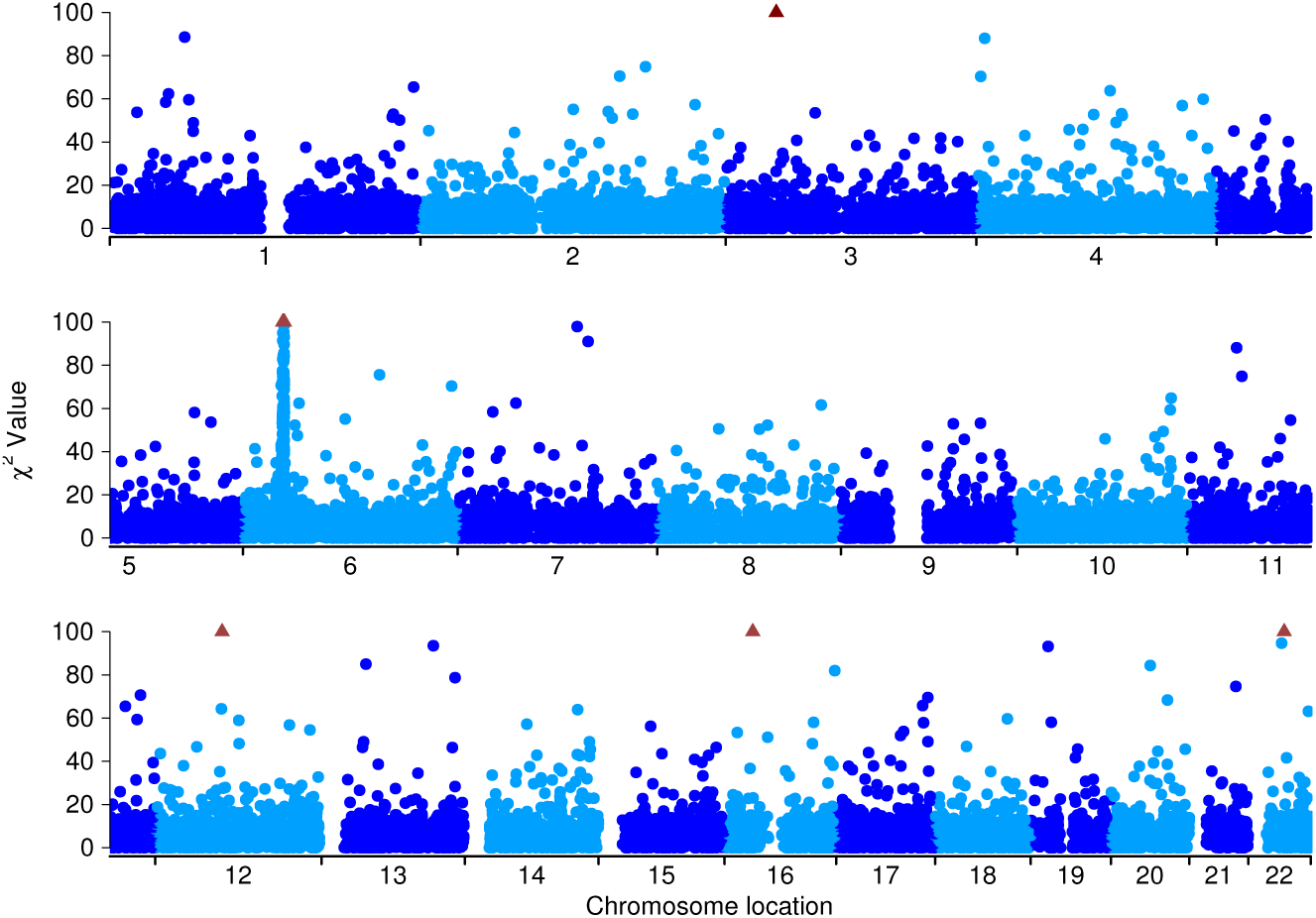
Scatter plot indicating unfiltered marker association values across the autosomal genome in the discovery group. Values greater than the range of this graph (*χ*^2^ *>* 100) are shown at the top of the graphs as a triangle symbol. A large spike of high association values can be seen near the middle of the short arm of chromosome 6.

The bootstrap sub-sampling method was used to eliminate those markers from the initial X chromosome filtered set of 490032 SNPs that were not effective for genetically distinguishing case and control groups. In each iteration of the bootstrap process, a sub-sample of individuals from each group was carried out, then markers were ranked based on a statistic that evaluates the effectiveness of each marker (see Figure 4). Markers that consistently had a high association statistic in each bootstrap sub-sample were selected for the next stage in the process.

#### 2.3.1 Comparison of Bootstrap Sub-sampling With a Simple Ranking Method

The bootstrap sub-sampling process attempts to eliminate markers that are specific to the particular sample of individuals under study, rather than the more general population those individuals have been sampled from. The effect of using a simple ranking procedure that has no sub-sampling would be to identify the markers that are most differentiated in that particular sample of individuals. However, natural variation in genotype frequency introduces noise into association analyses, so markers that are differentiated in a particular sample may not be differentiated in the population the sample was derived from.

The problem of discovering associated features that are not present in the more general case is known as overfitting [see ? , Chapter 14, pp. 661-663]. In the conventional GWAS context, overfitting produces false positive associations, where an association with a particular genotype does not extend to the general population. Using a method that includes bootstrap sub-sampling should reduce the degree of overfitting by removing markers that are only relevant for distinguishing between the specific groups involved in marker discovery.

**Figure 3:**
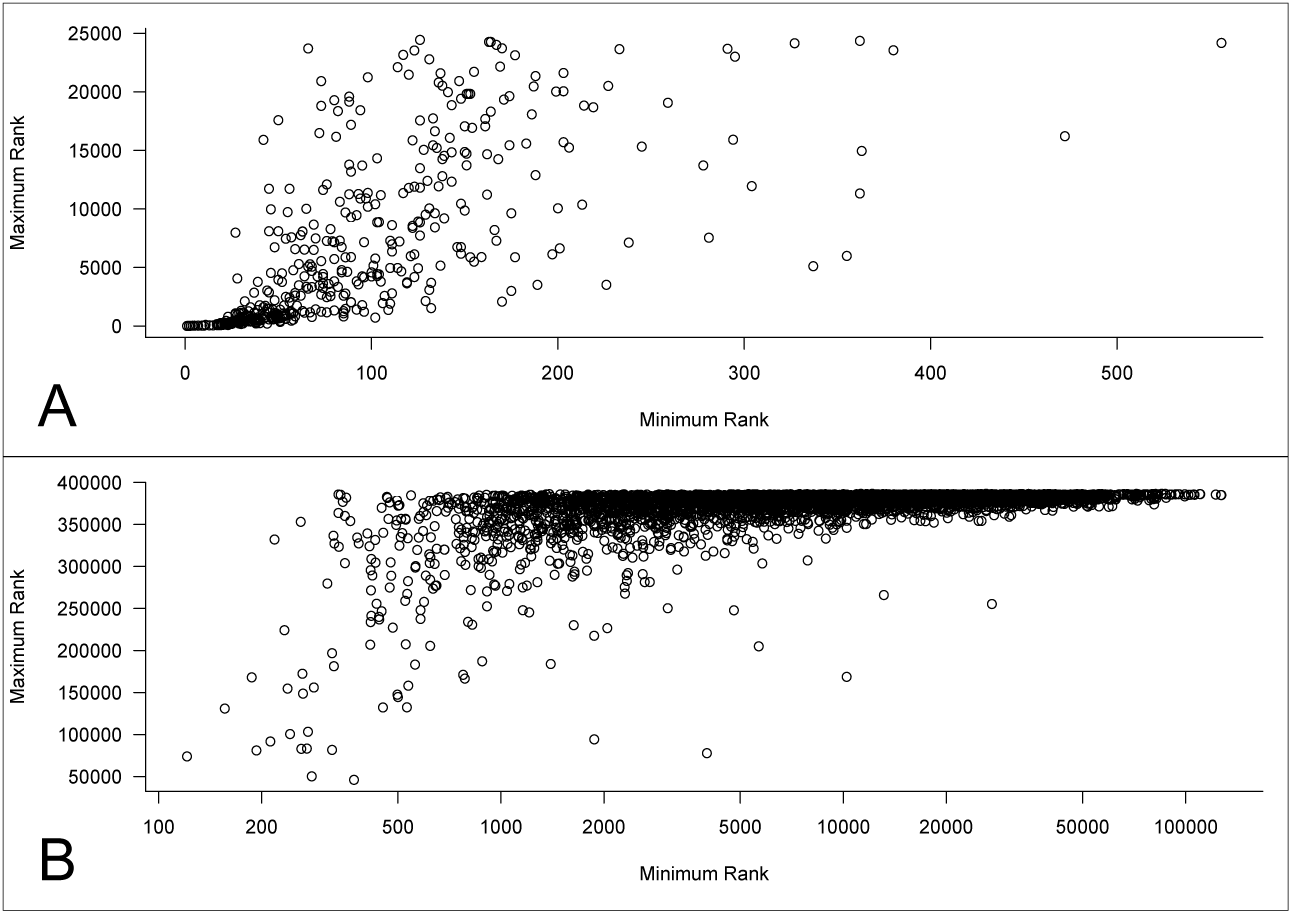
Scatter plot indicating maximum rank over all bootstrap sub-samples vs. minimum rank in any bootstrap sub-sample for the bootstrap-consistent set of 458 SNPs (A), and a random sample of 5000 of the remaining SNPs (B). A total of 126519 SNPs (not included when generating these graphs) were unranked in at least one bootstrap sub-sample, as no genetic difference was observed between case and control groups with that SNP. The difference between minimum and maximum rank gives an indication of the reliability of a particular marker for association testing in a general population. Of those markers in the bootstrap-consistent set of 458 SNPs, 57% were ranked in the top 5000 markers in all bootstraps. Of the remaining 489574 SNPs, 95% (4464977) had a maximum rank of 350000 or more (including 126519 unranked SNPs).

#### 2.3.2 Choosing a Marker Ranking Statistic

A ranking statistic is necessary for the bootstrap process to determine which markers are more likely to be associated with the phenotype of interest.

The purpose of the ranking statistic is to rank the effectiveness of markers in distinguishing groups, rather than give a precise indication of their utility. This means that the actual statistic used is not important, as long as it is generally able to rank an informative marker higher than a less informative marker. In this case, a genotype-based *χ*^2^ statistic was chosen for evaluating marker effectiveness. This statistic considers situations where a heterozygous genotype may have a strong association that is not present in either homozygous genotype, as well as identifying strong associations for homozygous genotypes.

#### 2.3.3 Ranking Markers Using the Observed Distribution of Ranking Statistics

A non-parametric ranking method selected markers based on rank order across all bootstraps. Markers are assigned a rank within each bootstrap: the marker with the most informative statistic is assigned rank 1, the second-most informative is assigned rank 2, and so on. The minimum, maximum, and mean marker rank are determined for each marker in all bootstrap sub-samples (see Table 1).

#### 2.3.4 Identifying Group-specific Markers

Markers that are not ranked in the top 5% of markers in *any* sub-sample are excluded from further analysis. When using this process on the T1D discovery group, a bootstrap-consistent set of 458 group-specific markers (GSMs) were found in the top 24501^3^ markers in *all* 100 sub-samples. Of these GSMs, 182 (40%) are located between 30Mb and 33Mb from the beginning of chromosome 6, near the HLA region. The remaining 276 GSMs are distributed fairly evenly throughout the genome (see Figure 5). From these observations of chromosomal location, T1D appears to have a very strong association signal near the HLA region on chromosome 6, and limited signal elsewhere in the genome.

**Table 1:** A sample of markers from the T1D study, showing minimum, maximum, and mean rank in 100 bootstrap sub-samples. In order to demonstrate differences between included (low rank in *all* bootstrap sub-samples) and excluded markers, the first ten markers were sampled randomly from the group of 458 markers with maximum rank less than 24502, and the remaining ten markers were sampled randomly from the remaining 489574 markers.

| Marker | Min Rank | Max Rank | Mean Rank |
| --- | --- | --- | --- |
| rs2027852 | 9 | 27 | 17.8 |
| rs3135342 | 42 | 1598 | 196.2 |
| rs16917773 | 48 | 2196 | 261.5 |
| rs10144861 | 59 | 2817 | 491.3 |
| rs10842028 | 44 | 3028 | 543.4 |
| rs10742084 | 61 | 5290 | 679.9 |
| rs7158350 | 84 | 6736 | 730.4 |
| rs16854531 | 126 | 13477 | 735.1 |
| rs1429445 | 54 | 7451 | 975.9 |
| rs17023486 | 472 | 16210 | 2903.6 |
| rs12117563 | 1508 | 377782 | 77651.8 |
| rs11189528 | 1563 | 385065 | 162459.9 |
| rs17038075 | 47938 | 453037 | 172338.7 |
| rs11081211 | 13893 | 376453 | 185579.0 |
| rs11249611 | 1731 | 376943 | 190683.0 |
| rs16831752 | 119403 | 461199 | 222291.4 |
| rs1006931 | 22398 | 369366 | 232357.9 |
| rs9317562 | 15117 | 383671 | 235382.3 |
| rs11205709 | 477789 | 477983 | 477890.9 |
| rs1027341 | 487082 | 487165 | 487114.9 |

**Figure 4:**
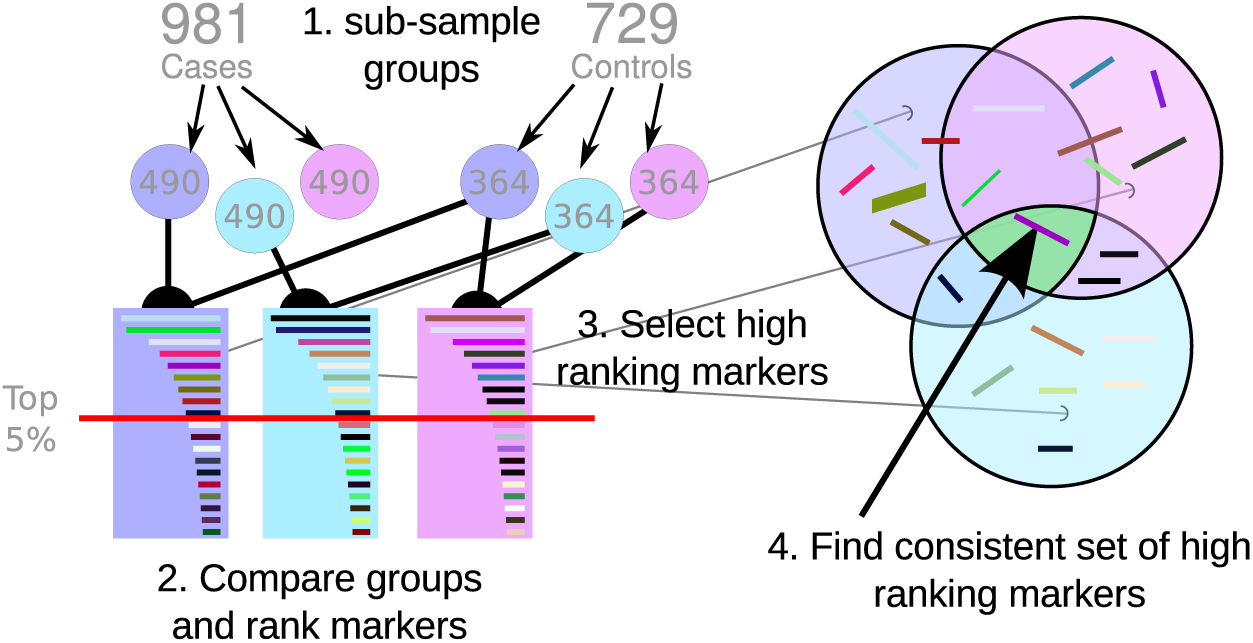
Visual representation of the key points of the bootstrap process. The groups are sub-sampled a number of times, and marker ranking statistics are calculated for each sub-sample (bootstrap). Markers are then ranked, identifying the markers with the highest association statistic for each sub-sample. Markers that were consistently ranked in the top 5% in all sub-samples were passed onto the next stage of the selection process.

### 2.4 Linkage Refinement

Linked GSMs were removed in order to reduce the redundancy of associative signal produced by the generated GSM set. Markers were ordered based on mean rank order and any SNPs that were linked (*r*^2^ *>* 0.1) with a higher-ranked GSM were removed from the set, leaving an *unlinked set* of 34 GSMs.

Markers within a signature marker set should be unlinked, so it is a good idea to calculate a linkage-associated statistic such as *D'* or *r*^2^ during the discovery phase of the analysis, and remove the least informative marker among linked high-association pairs. This step is carried out after the bootstrap sub-sampling process in order to reduce the number of pairwise calculations required for linkage analysis – pairwise calculations for 500 markers would require 124,750 linkage comparisons,^4^ while pairwise calculations on 500,000 markers would require around 1.25 *×* 10^11^ comparisons.

### 2.5 Set Size Refinement

The optimal marker set size was identified using an Area Under the Curve (AUC) test on the Q-values generated by *structure* (10,000 bootstraps, and 100,000 total runs), finding marker sets with large differences in mean Q value between the two groups (see Figure 6). Increasing numbers of markers were selected from the unlinked GSM set based on mean rank order identified during the previous (bootstrap sub-sampling) stage. The effectiveness of a given set of markers was evaluated using the *structure* program, followed by an AUC calculation for each set of markers based on Q values reported by the program.

**Figure 5:**
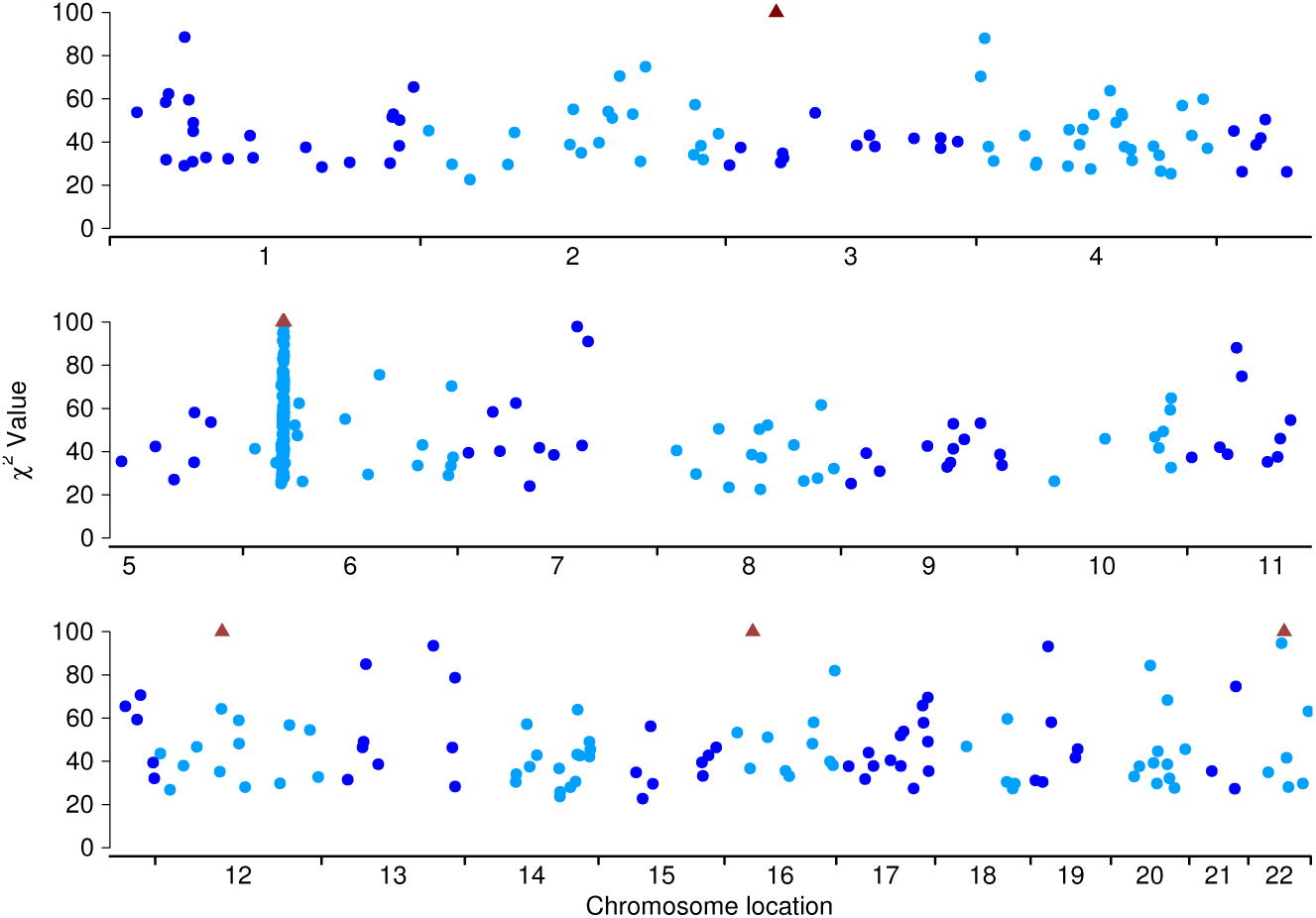
Scatter plot indicating marker association values for the consistent set of 458 GSMs across the autosomal genome in the discovery group. Values greater than the range of this graph (*χ*^2^ *>* 100) are shown at the top of the graphs as a triangle symbol. The large spike of high association values still remains near the middle of the short arm of chromosome 6.

The *structure* program outputs values that represent to how genetically similar an individual is to a particular group (Q values), attempting to cluster pooled individuals into two “populations”.^5^ The Q values produced by *structure* are continuous in the range between 0 and 1 inclusive, and are treated as an estimate of the probability that an individual has a particular trait.

Analysis of Q values was used to determine false positive and true positive rates for given Q-value cutoffs (see Figure 8). The true positive rate was calculated as the proportion of T1D cases with Q below the cutoff value, and false positive rate was calculated in the same way for NBS controls. The area under the curve of this graph can be used as an indication of the effectiveness of a quantitative test. An AUC of 1 indicates a perfect test (no misclassification), while an AUC of 0.5 indicates a test that cannot distinguish between groups.

The greatest difference between cases and controls was observed when the top 5 GSMs were selected, producing an AUC of 0.8449. This *signature set* of 5 GSMs was considered to be the most appropriate T1D-informative set.

### 2.6 Validation of Final 5 GSM Set

The signature set of 5 GSMs (see Table 2) was finally tested on the validation group (982 T1D cases, 729 NBS controls) using *structure*, followed by an AUC analysis of the Q values. There is a small overlap between some T1D cases and some NBS controls (Figure 7), but most T1D cases cluster together, and are separate from the cluster of NBS controls.

The AUC value associated with this test of the signature set of 5 GSMs in the validation group was 0.8395. Setting the false positive rate to 5% (cutoff Q value 0.129) produced a true positive rate of 43%, while setting the true positive rate to 85% (cutoff Q value 0.5583) produced a false positive rate of 38%. The position on the curve nearest to a true positive rate of 100% and a false positive rate of 0% was when the cutoff Q value was set at 0.506, with a true positive rate of 78%, and a false positive rate of 29%.

**Table 2:** Location information for the top 5 GSMs discovered in a bootstrap sub-sampled GWAS for T1D associations, after removing linked GSMs, and choosing the set with the highest AUC value. Mean rank reported in this table is based on the marker rank for 100 bootstrap sub-samples. Out of the five markers, four are within a 2Mb region of chromosome 6.

| Marker | Chromosome | Location (Mb) | $\chi^2$ | Mean Rank |
| --- | --- | --- | --- | --- |
| rs9273363 | 6 | 32734250 | 485 | 1 |
| rs3957146 | 6 | 32789508 | 317 | 2.2 |
| rs3135377 | 6 | 32493377 | 264 | 4.3 |
| rs7431934 | 3 | 40268801 | 199 | 13.7 |
| rs1046089 | 6 | 31710946 | 108 | 37.9 |

**Figure 6:**
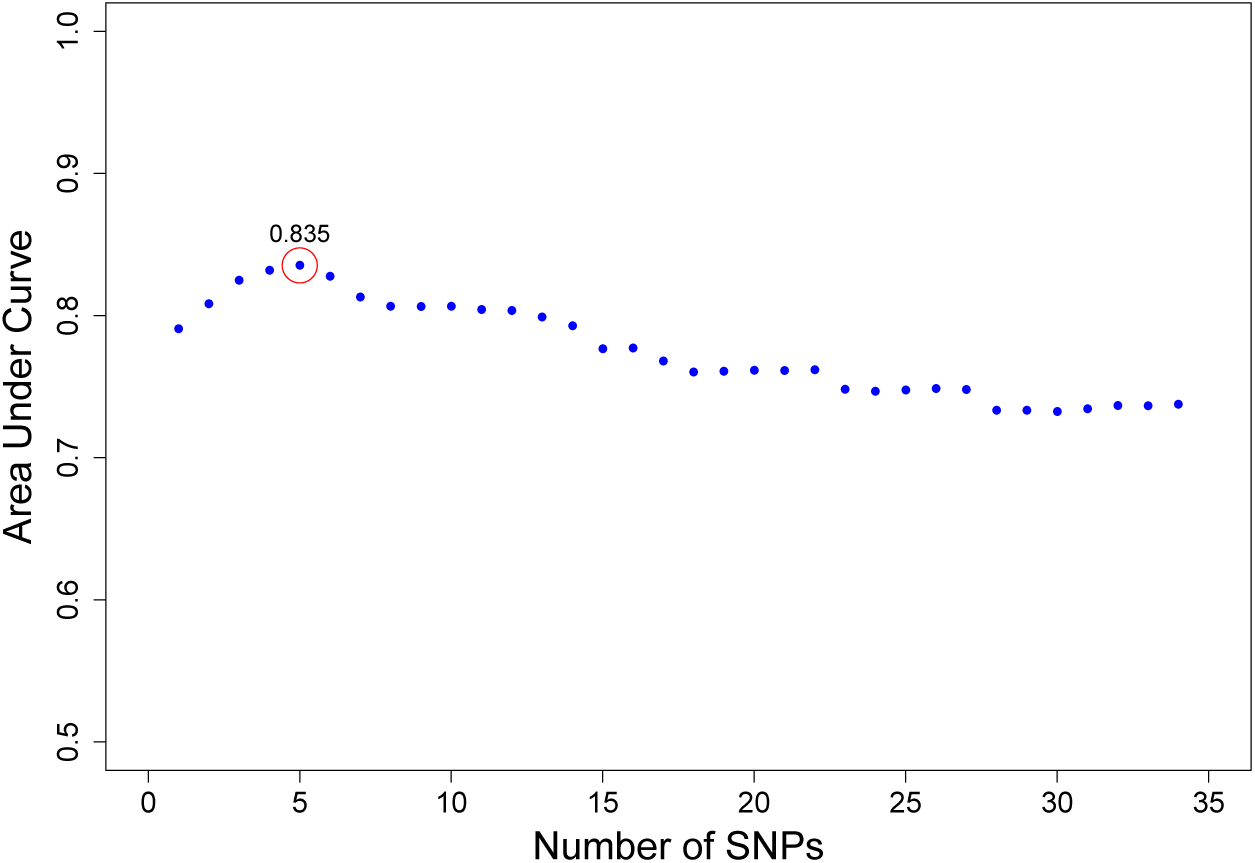
A marker refinement plot, showing the effectiveness score (AUC) for increasing numbers of SNPs in the discovery group. The highest AUC value (0.835 for 5 SNPs) is circled in red.

### 2.7 Comparison with SNP set from Literature

Todd et al. [11] carried out an analysis of 11 SNPs that were found to be associated with Type 1 Diabetes in genome-wide association studies. This group of SNPs, in combination with the most informative SNP from the WTCCC study [12], was selected to be compared with the signature set of 5 GSMs in the present study (see Table 3). The *structure* program was used in combination with an AUC analysis to evaluate the effectiveness of this group of 12 SNPs for 1963 WTCCC T1D cases and 1458 NBS controls (see Figure 9 and Figure 10).

This 12-SNP comparison set had an AUC of 0.73 when tested with 1963 T1D cases and 1458 NBS controls. Setting the false positive rate to 5% (cutoff Q value 0.933) produced a true positive rate of 18%, while setting the true positive rate to 85% (cutoff Q value 0.895) produced a false positive rate of 53%. The position on the curve nearest to a true positive rate of 100% and a false positive rate of 0% was when the cutoff Q value was set at 0.910, with a true positive rate of 65%, and a false positive rate of 35%. These results indicate that the signature GSM set discovered in the present study is considerably more informative than a set of T1D-associated SNPs found in other genome-wide association studies.

**Figure 7:**
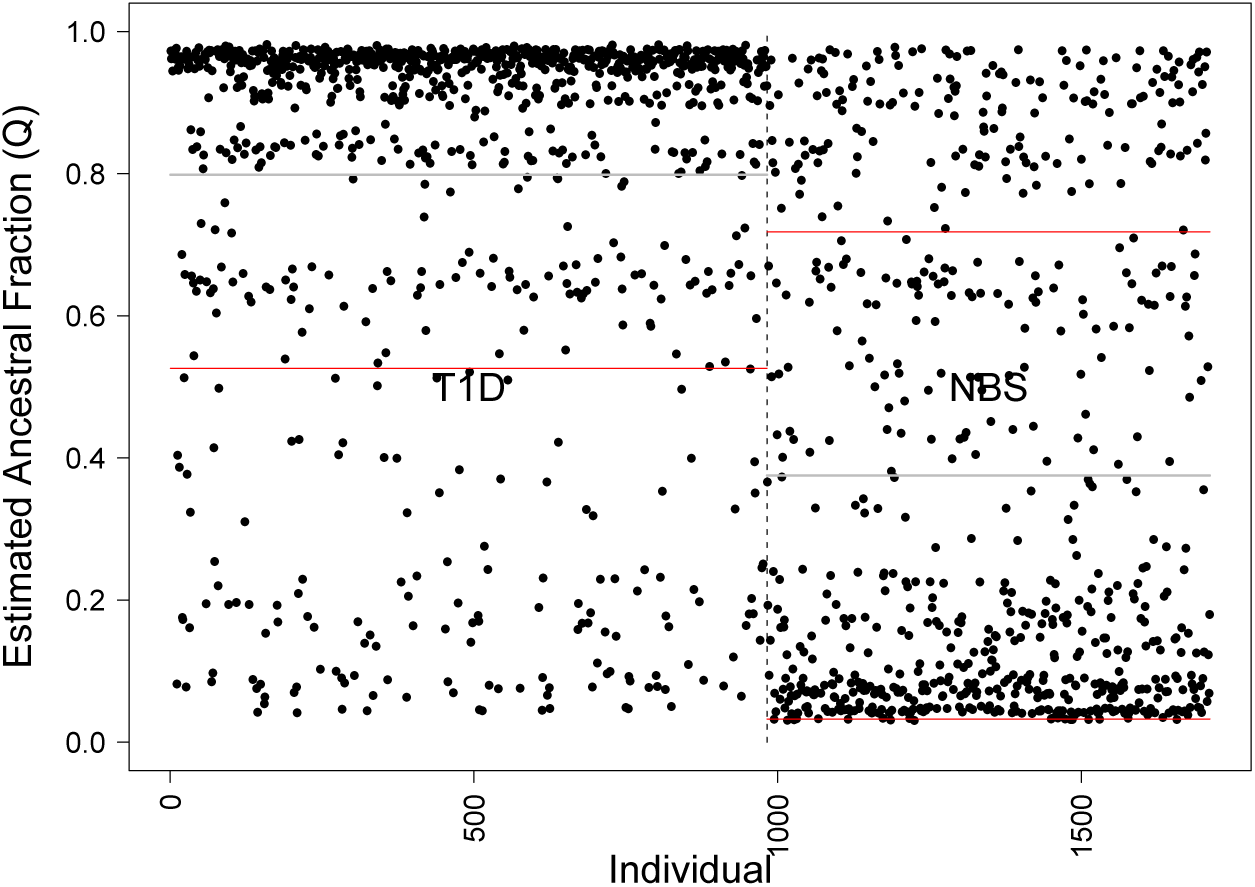
Structure output (K=2) for the top 5 SNPs discovered in a bootstrap sub-sampled GWAS for T1D associations, showing Q values for individuals from T1D and NBS groups (using validated group).

## 3 Discussion

This study has identified a group of 5 GSMs that classify individuals with T1D with good reliability (AUC = 0.84, see Figure 8). The heritability of Type 1 Diabetes is around 88% [6], so the maximum possible sensitivity (true positive rate) of a genetic test for T1D should be 88%, with the remaining 12% of variation being due to non-genetic factors.

One of the assumptions made in GWAS is that the individuals selected as candidates for the phenotypic groups (cases and controls) are ideal members of those groups – affectation status tends to be a binary or integer value that does not allow for intermediate values. Due to the difficulty in qualitatively describing traits, as well as mutation and admixture effects (particularly for population-derived groups), this assumption may be invalidated.

The marker construction method used a bootstrapping procedure as an internal validation to remove markers that had substantial variation in *χ*^2^ values within the tested groups. In an ideal case, a bootstrapping procedure would not be necessary as the genetic makeup of the total population will reflect the makeup of any given subgroup of that population. In such a case, the ranking after each bootstrap will be the same as the overall ranking. However, the comparison of minimum and maximum rankings for SNPs across all bootstrap sub-samples has demonstrated that this is clearly not the case (see Section 2.3).

**Figure 8:**
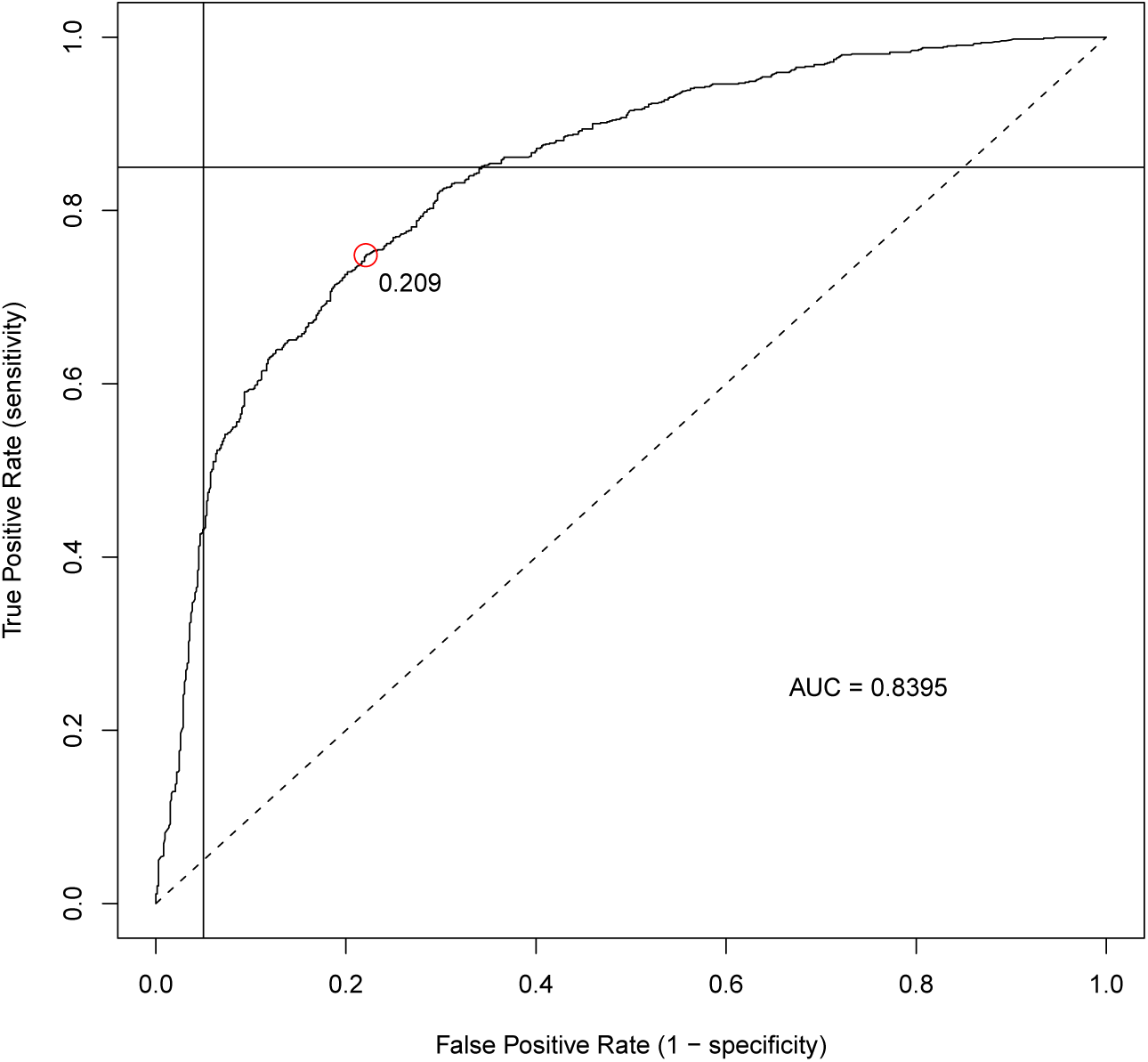
Line graph of true positive rate vs. false positive rate (Receiver-Operator Curve) based on structure plot of validated set of 5 SNPs.

### 3.1 Type 1 Diabetes Study Results

It is known that genetic variation within the HLA region on chromosome 6 plays an important role in T1D, accounting for about 50% of the genetic susceptibility for T1D [see 3]. This role is supported by the preliminary results in the present study, which show consistently strong predictive power using genetic markers, all but one from this region alone (see Table 2).

#### 3.1.1 Accuracy of the Signature SNP Set

The interpretation of accuracy of a genetic test is difficult, particularly when considering what would be expected if the test were used in an untested population. A statistic that can be useful in this case is the positive predictive value (how likely a test is positive, given a positive result).

In order to determine the positive predictive value of a test, it is necessary to establish the prevalence of the trait in the population of individuals who are to be tested. A country which is considered to have a very high incidence of T1D, Finland, has an overall cumulative incidence of around 0.5-0.6% at the age of 35 years [6]. Also, there has been a general trend of a 2-3% increase in the incidence rate of childhood T1D in South West England over the past 20-30 years, with the incidence in 2003 at around 0.16% per year [13]. Even at the higher incidence rate in Finland, fewer than 0.6% of individuals in a typical non-enriched control population would be expected to have T1D.

**Table 3:**
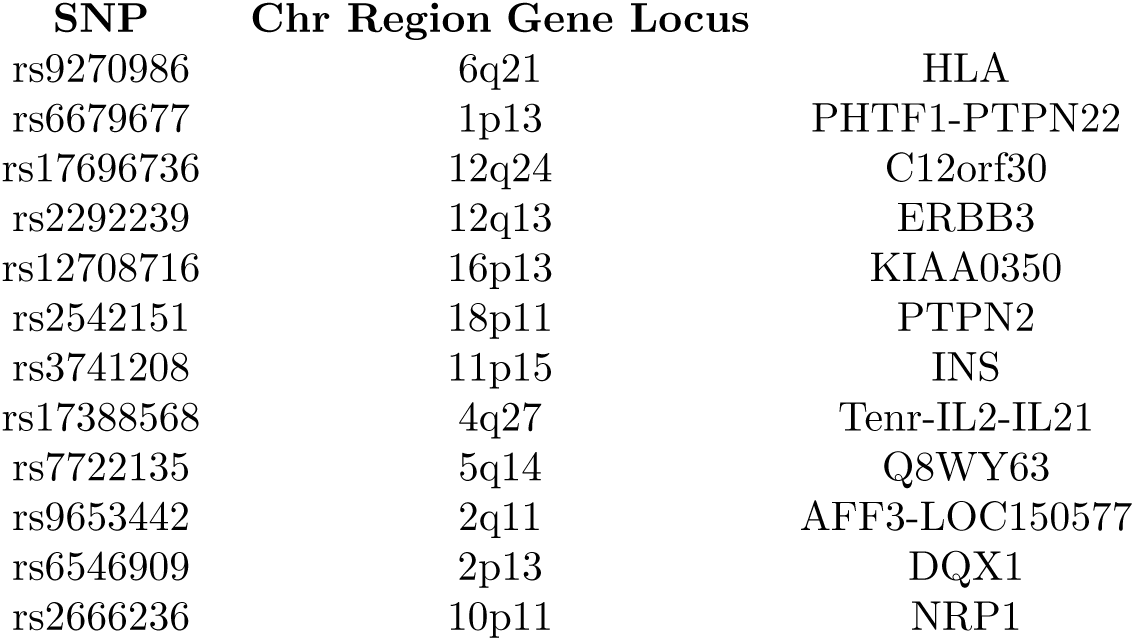
A list of SNPs found by other researchers to be associated with T1D risk. The first SNP (rs9270986) yielded the most extreme statistic in the WTCCC analysis [12]. Marker names and locations for the remaining 11 SNPs are from Table 1 of Todd et al. [11].

**Figure 9:**
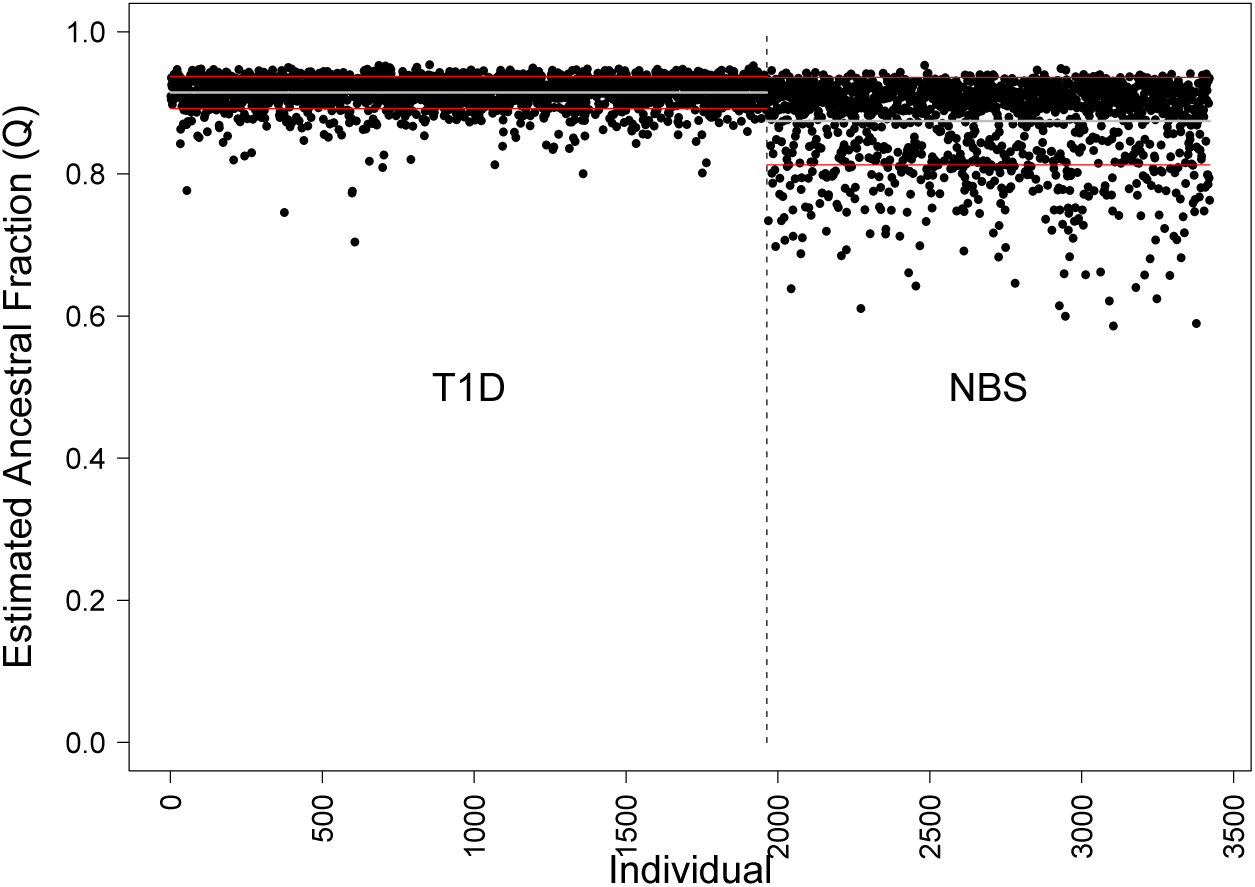
Structure output (K=2) for 12 SNPs found by other researchers to be associated with T1D risk (see Table 2).

The NBS controls for the WTCCC study had not been enriched to remove individuals that have T1D. Given an expected prevalence of T1D of 0.6%, it would be expected that around 4 individuals from the validation NBS control group (or 9 from the discovery and validation groups combined) have T1D. Setting the false positive error rate to this value (i.e. 0.6%) is unrealistic for the current data set, as only a small fraction of T1D cases would be identified with that cutoff (just over 5%, see Figure 8). However, if a more moderate 5% false positive error rate is accepted (identifying 43% of T1D cases, see Section 2.6), then 36 NBS individuals would be identified by this test as at risk for T1D. This is about ten times that expected by cumulative incidence rates for T1D, indicating a positive predictive value of 10% with the discovered signature set of 5 GSMs. Given that the population prevalence of T1D is so low, the NBS control group should not differ substantially from an enriched control group, and the positive predictive value of this genetic test will remain around 10%.

**Figure 10:**
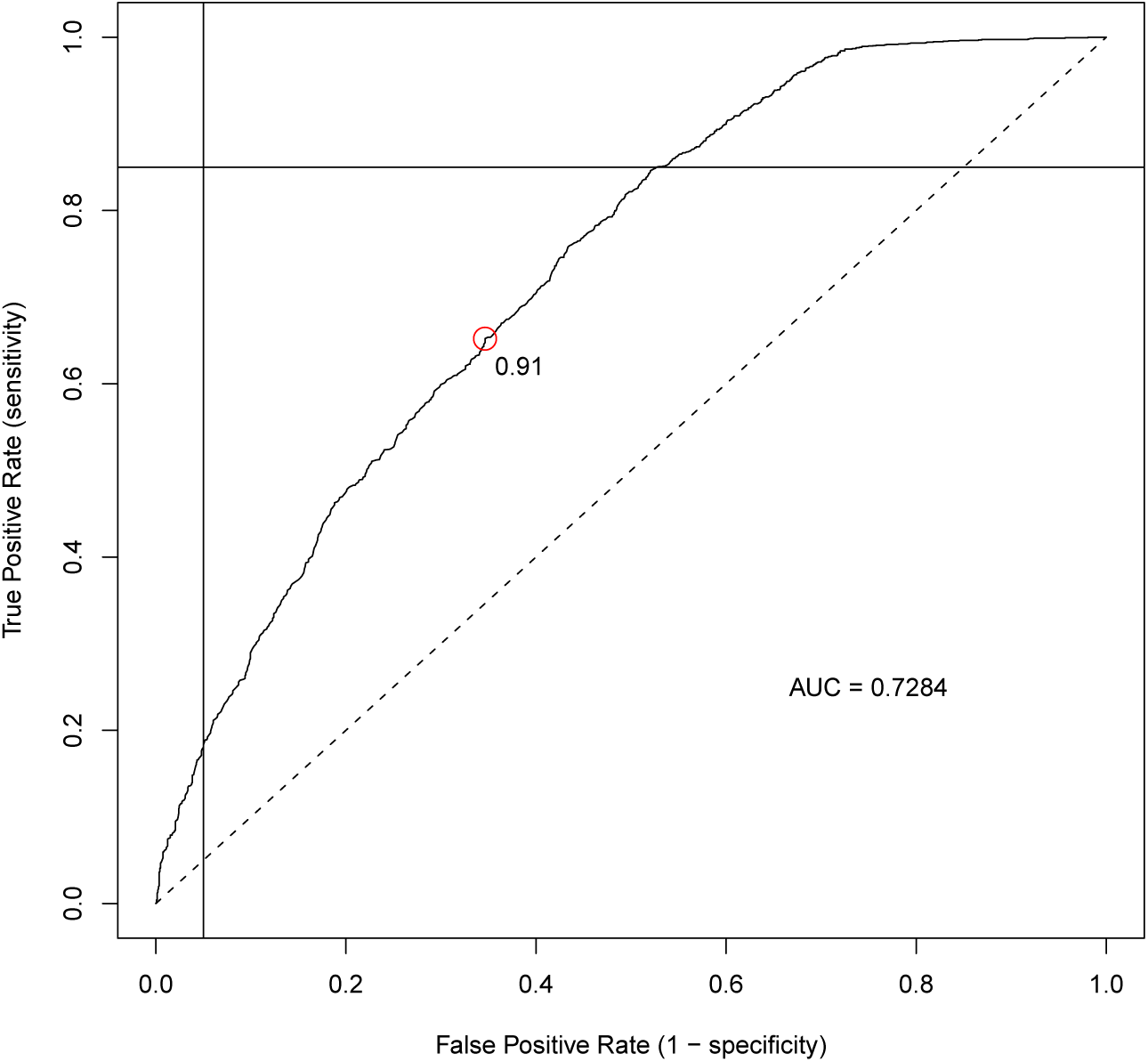
Line graph of true positive rate vs. false positive rate based on structure plot of SNPs found by other researchers (see Table 2).

However, if it is the case that a substantial proportion of Type 2 Diabetes cases are actually Type 1 Diabetes, there may be an underestimation of of T1D incidence in the general population [see 1]. Such a disparity would substantially increase the predictive value of a genetic test using the GSMs discovered here.

#### 3.1.2 Accuracy in Other Populations

The low positive predictive value of the marker set, together with heritability values of less than 100%, means that it is unlikely that a genetic test using these T1D markers would be useful as a *diagnostic* test for a general population. However, if used in conjunction with other clinical indicators, it may be appropriate to use these genetic markers for a *screening* test, identifying individuals that should be more closely monitored for T1D symptoms. This is because it will still exclude a large proportion of the normal population, while also identifying a high proportion of at-risk individuals. However, the signature GSM set has not been validated in groups of individuals outside the WTCCC study, and caution should be taken in attempting to extrapolate results to non-validated populations.

Taken in the context of disease, it can be very difficult to accurately determine the phenotype of an individual – this is a particular problem when the disease is a continuous (rather than discrete) trait, as often happens with common complex diseases. Phenotype identification is further complicated by non-Mendelian patterns of inheritance. It is possible for there to be numerous paths to the same apparent end disease, and numerous gene-gene interactions that contribute to the same disease. Furthermore, trait variation is often a mixture of genetic and environmental factors (i.e. heritability is less than 100%), so potential gene-environment interactions also need to be taken into account when describing phenotype.

The effectiveness of any given set of markers will be reduced due to the presence of erroneous false positive results (i.e. some of the false positives will later turn out to have T1D). In a situation where the marker set is constructed to remove as many false positive results as possible, this may result in a refined test that is over-fitted to the initial discovery group of case and control individuals, and is not reliably generalisable to other populations. It is possible that such situations would be apparent when follow-up studies on independent case/control groups for the same trait are carried out, and it is recommended that such validations are carried out before using this signature GSM set.

### 3.2 Overfitting Generates Spurious Associations

For a genetic association study to be successful, individuals must be separable into distinct groups based on a particular phenotype, and some differences between the groups must be attributable to genetic factors. Methods for identifying associated markers in a GWAS relies on a clear distinction between trait and non-trait individuals. In situations where the trait of interest is not easy to classify, an associated marker may not reflect the true distinction between those groups. In addition, a low genetic influence for the expression of a particular trait can mean that even when a trait can be classified completely, the genetic component of that trait (the only component able to be identified by any DNA marker-based method) will not always determine the observed phenotype completely.

Overfitting is the generation of a set of distinctive parameters that relies on irrelevant attributes for the model being observed. The problem exists when vital information about the model is missing, and the discovery algorithm ends up being required to derive a model based on other spurious distinctions between discovery groups [see ? , Chapter 14, pp. 661-663]. Overfitting is applicable to the case of generating minimal marker sets because any such method assumes that a minimal set can be found for the data. When cases and controls are not genetically distinct, and distinct *only* due to the trait under test, any resultant marker set will be invalid. In such a situation, the set of markers generated is informative only for the specific group of individuals that were used for discovery of that set of markers, and will not be applicable for individuals outside the discovery group. Internal validation within groups, and external validation of results in similar populations, is essential to ensure that overfitting has not occurred.

Bootstrap sub-sampling uses variance among group sub-samples to remove markers that are associated because of *genetic chance* effects rather than the particular phenotype under test. However, it cannot distinguish between genetic differences due to the tested phenotype and genetic differences due to sampling bias. The problem of overfitting is especially relevant for genetic data, where one pattern of genotypes due to a group-associated factor with high heritability may outweigh the disease-causing factor under test. This is similar to the population stratification problem that has been discussed by Pritchard [8] and Pritchard & Donnelly [9] who say that due to the influence of *genetic chance* (e.g. genetic drift, founder effects, non-random mating), alleles can appear with high frequency differences between groups within a given population sample even though the differences are not directly associated with the trait of interest. This is particularly important when a population group has a high incidence of a given disease, and the genetic history of the case and/or control subgroups is not known. Pritchard & Donnelly [9] recommend testing for structured association in case and control groups before carrying out further association tests in order to remove confounding genetic factors that may be present in a case/control study.

#### 3.2.1 Genome-wide Trait Contributions

While there may be many gene-gene interactions throughout the genome that all contribute to a particular disease, it is unlikely that *all* genetic variants in the subgroup will influence the trait. In addition, some variants may influence the trait more than others and in some cases may even negate the effects of another variant. Both of these factors increase the potential for spurious associations and false positive results when carrying out a whole genome scan.

Genotyping carried out in an association study is restricted to a subset of the total genome, because full-genome sequencing is still prohibitively expensive. Also, only a subset of interactions between multiple genetic factors can be studied (if any), because multi-factorial analysis is computationally expensive.^6^

It is expected that any reduction of GSM set size will result in decreased reliability, as there is an information loss when fewer markers are typed. For a reduction method to be useful, the information lost due to typing fewer markers must be compensated by cost reduction. However, in this investigation, the opposite appears to be true – a small number of markers are useful to distinguish the case and control groups, and appear to provide more information than a full genome set.

#### 3.2.2 Interactions from Multiple Genetic Variants

In some cases, a first-pass single association analysis of markers will not be useful for the classification of a trait. This will be the case for traits that have complex interactions that result in non-linear association patterns between marker frequency and trait prevalence. As an example of a complex interaction, two causative variants may interact in a neutralising fashion (i.e. the effects of one variant are cancelled out by another variant). In this sort of case, a simple one-way association test would not work as expected, retaining a lack of observed association even when there is a strong signal [7]. Other non-linear interactions between different markers would also reduce the effectiveness of an association test to determine informative markers.

The ideal situation for investigating complex traits at a genetic level is an analysis of the effectiveness of *every possible* set of marker interactions. Once such an analysis is carried out, the best set of markers will be identified as being the set that is most informative for classifying individuals into groups. However, the computational requirements for such testing combined with the increased danger of overfitting due to small cell sizes, make such an analysis effectively useless when carried out on the total marker set [see 10].

The bootstrapping approach as outlined here does not consider combinations of genetic markers. However, it provides an efficient way to reduce a large set of markers down to a much smaller set. This smaller set can then be used by programs that determine multi-way interactions, which are typically very computationally expensive procedures.

## 4 Conclusion

The application of the bootstrap sub-sampling process to marker selection is a useful complement to current GWAS. It can be used to remove potential spurious associations that are specific to the tested groups, and may help to reduce the set of individuals required for initial large-scale genotyping. Bootstrap sub-sampling acts as an internal validation of association signals, which helps to reduce the likelihood of false positive associations in publications. This, in turn, would hopefully make clinicians less likely to use these false positive associations when evaluating disease risk.

The method for identifying a minimal set of GSMs is an association-based method that discovers genome-wide combinations of markers for the identification of a particular trait. The method relies on a clear distinction between trait and non-trait individuals, and in situations where the trait of interest is not easy to qualify, the identified GSM set may not reflect the true distinction between those groups. In addition, the non-genetic influence for the expression of a particular trait can mean that even when a trait can be classified completely, the genetic component of that trait (the only component able to be identified by any DNA marker-based method) will not always determine the observed phenotype.

It is essential that this set be externally validated in other populations, but given reasonable validation the set may also be used for a global indicator of T1D risk. Even without such validation, the Wellcome Trust study was a fairly large study that recruited a substantial proportion of UK individuals. Given this, the signature set of 5 GSMs identified here should be suitable for estimating T1D risk for screening purposes in at least the UK population.

1 i.e. taken at any time of the day, as opposed to a *fasting* glucose test taken at least 8 hours after the last meal

2 The study also typed 2000 individuals for each of the six other diseases: a total of 14,000 cases genotyped for seven diseases.

3 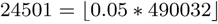

4 124, 750 = (500^2^ *−* 500)/2

5 The *structure* program is designed for *population* analysis, but is used here for *group* analysis.

6 It has an exponential complexity with respect to the number of factors studied in tandem.

## 5 Author Contributions

DAE carried out the data analysis and wrote the paper. GKC and RAL provided academic support for the research project, as well as editorial advice for the paper.

